# Structural basis of lenacapavir-induced HIV-1 capsid disruption during virion maturation

**DOI:** 10.64898/2026.02.16.706074

**Authors:** Hiroki Tanaka, Reina Morita, Tomomasa Oka, Shunsuke Kita, Mina Sasaki, Katsumi Maenaka, Shinichi Machida

## Abstract

Long-acting lenacapavir (LEN) has emerged as a highly effective, potentially game-changing therapy for HIV treatment and prevention^1–5^. Although its mechanism of action in the early phase of HIV-1 replication, when the capsid directs key post-entry steps such as reverse transcription, nuclear import, and integration, has been well characterized^1,2,6–8^, its effects during the late phase of replication, when the capsid assembles and matures within budding virions, remain poorly understood. Here, we determine the cryo-electron microscopy structure of the mature HIV-1 capsid lattice assembled within virus-like particles in the presence of LEN. Our structural analyses revealed that LEN alters interhexamer interactions, perturbs capsid lattice curvature, and thereby prevents the formation of a functional cone-shaped capsid. Biochemical analyses further demonstrated that LEN-containing cores lose reverse transcriptase due to a compromised capsid integrity, whereas integrase and viral RNA remain associated. Functionally, viruses produced in the presence of LEN exhibited markedly reduced infectivity, low reverse transcription activity, and poor integration. Together, these findings provide mechanistic insight into the late-phase action of LEN and provide key directions for the design of future inhibitors.

## Introduction

The HIV-1 capsid is a multifunctional structural protein complex that not only encloses the viral RNA genome but also regulates reverse transcription, nuclear import, and integration site selection, as well as evasion of immune sensing by cytosolic DNA sensors^9–13^. The HIV-1 capsid is composed of capsid protein (CA), which is released following the cleavage of the Gag polyprotein during viral maturation^14–16^. CA contains two distinct domains, the N-terminal domain (NTD) and the C-terminal domain (CTD), connected by a flexible linker. In mature virions, CA monomers assemble into a fullerene-like cone in which approximately 200 CA hexamers and 12 CA pentamers are arranged in a continuous lattice to form the conical capsid ^17–21^. The CA-NTD stabilizes each hexamer through interactions around the sixfold axis, whereas the CA-CTD mediates dimeric and trimeric contacts, involving helix 9 and primarily helix 10 of the CTD, respectively, to interconnect hexamers and build the lattice ^21–23^. Inositol hexakisphosphate (IP6) further stabilizes the lattice by binding to a ring of arginine residues within the CA-NTD, thereby promoting proper capsid assembly^24,25^. The precise balance between the stability and instability of the capsid is critical for its function during the viral life cycle^26–30^. Accordingly, the capsid represents a pivotal target for HIV-1 therapeutic intervention.

Lenacapavir (LEN, also known as GS-6207) is a first-in-class HIV capsid inhibitor that exhibits high potency and long-acting activity, which allows it to be administered as infrequently as twice per year^3,4^. Furthermore, preexposure prophylaxis (PrEP) with LEN administered twice a year has been shown to markedly reduce the risk of HIV acquisition^5^. LEN has been shown to inhibit multiple steps of the viral replication cycle. Previous studies have demonstrated that LEN and GS-CA1, an archetypal predecessor of LEN, target the mature HIV-1 capsid and bind to the FG (phenylalanine–glycine motif)-binding pocket located at the interface between the NTD and the CTD of CA monomers, thereby altering the stability of the mature capsid, potentially modulating host factor binding to CA and inhibiting reverse transcription, nuclear import, and integration ^1,2,6–8^. Although the inhibitory effects of LEN have been characterized primarily in the context of postentry processes, its mechanistic impact during the late phase of HIV-1 replication, when the capsid assembles and matures within budding virions, remains poorly understood. Notably, LEN also binds to Gag monomers with high affinity, leading to the formation of aberrant capsid in virions and a marked reduction in viral infectivity^1,6^. In addition, a previous study suggested that LEN preferentially stabilizes the hexameric dimer interface over the pentameric interface, thereby biasing assembly towards hexamer formation^31^. Furthermore, the previous structural study revealed LEN binds to the immature HIV-1 gag lattice ^32^.

Despite these studies, it remains unclear how LEN-bound capsids reorganize during virion assembly or whether LEN incorporation alters the curvature parameters required for proper cone closure. Thus, defining how LEN reshapes capsid curvature during virion maturation is essential to fully understand its antiviral mechanism.

### LEN perturbs capsid lattice curvature in mature HIV-1 virus-like particles

To investigate the mechanistic effects of LEN during the late phase of HIV-1 replication, we examined the *in situ* structure of the HIV-1 capsid lattice within Gag virus-like particles (VLPs) produced in the presence of LEN. The VLPs produced from 293T cells in the absence or presence of LEN were vitrified and imaged by cryo-EM (Fig. 1a–c). Consistent with the findings of previous studies^1,6^, the VLPs produced in the presence of LEN predominantly contained irregularly shaped capsids rather than typical cone-shaped capsids (Fig. 1a–c). Interestingly, VLPs produced in the presence of LEN also showed a broader size distribution, and larger particles appeared more frequently in the presence of LEN than in its absence (Fig. 1a–c, Fig. S1). To obtain a higher-resolution view of the lattice architecture, we next treated the VLPs with perfringolysin O (PFO). This selectively perforated the viral membrane with a limited number of ~20–30 nm pores, thereby releasing excess free CA and increasing the signal-to-noise ratio in the cryo-EM images while maintaining overall membrane integrity and preserving the internal capsid structure in an apparently unaltered state^30,33,34^. We then determined the capsid lattice structure within VLPs produced in the presence of LEN using a single-particle cryo-EM approach (Fig. 1d, Figs. S2-3, Table S1). The reconstruction revealed a sixfold symmetric structure of the hexameric CA lattice within the virion capsid with LEN, in which the central CA hexamer was clearly resolved (Fig. 1e) and neighbouring hexamers were also discernible (Fig. 1d). A distinct density corresponding to LEN was observed at the expected FG binding pocket (Fig. 1f, g).

**Fig. 1.**
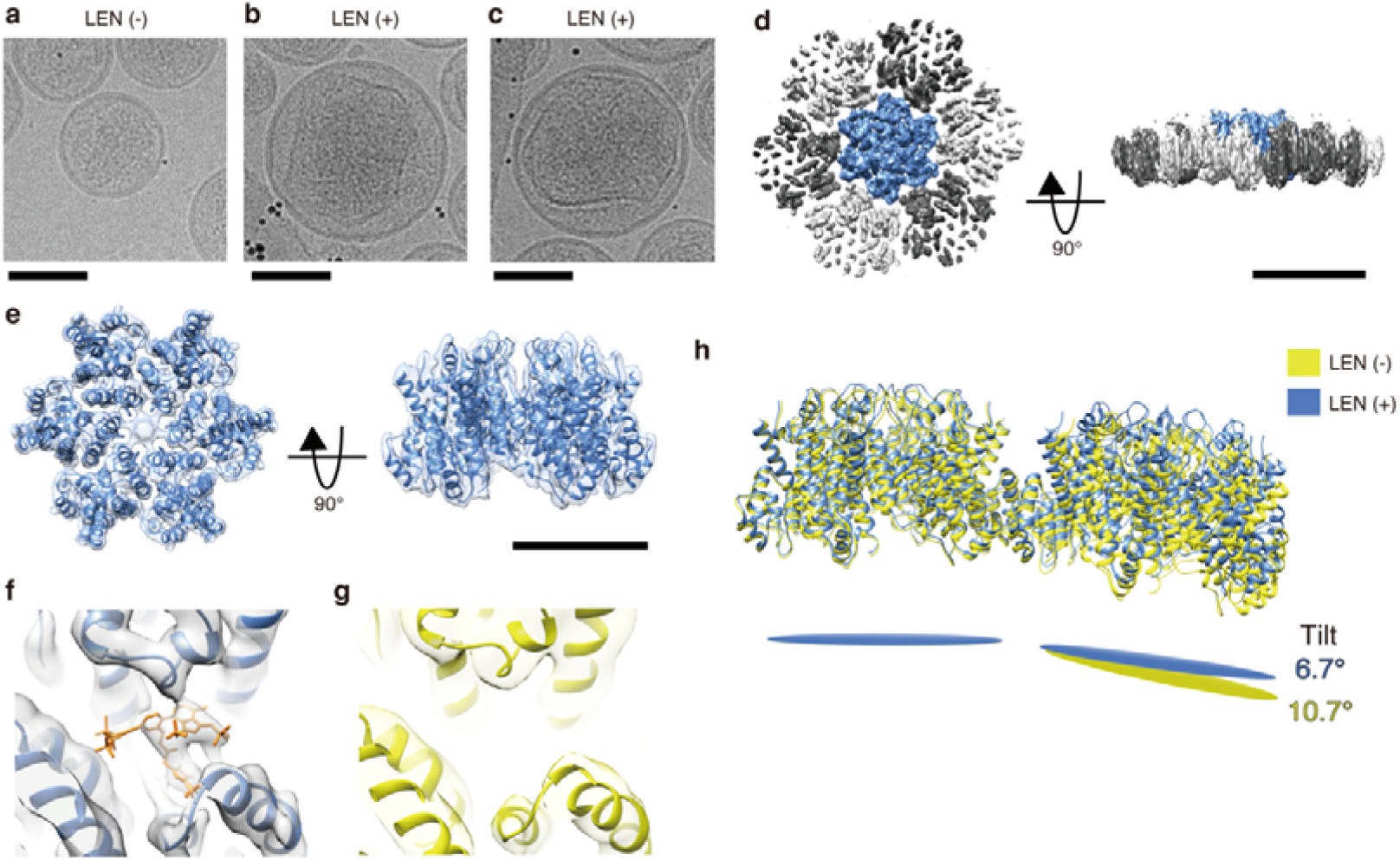
Cryo-EM structure of the HIV-1 capsid lattice within the VLPs produced in the presence of LEN. **a–c**, Cryo-EM images of the VLPs produced in the absence (a) or presence (b, c) of LEN. Scale bar, 100 nm. **d**, Cryo-EM density map of the CA lattice within the LEN-containing VLPs, shown from the top (left) and side (right) views. Scale bar, 100 Å. **e**, The atomic model of a CA hexamer in the LEN-bound lattice fitted into the cryo-EM density map shown in (d). Scale bar, 50 Å. **f**, Close-up view of the cryo-EM map of a LEN-bound CA lattice (d) and the atomic model (e) showing the density corresponding to a bound LEN. **g**, Close-up view of the cryo-ET map of a LEN-free CA lattice (EMDB: 13423) and its atomic model, showing the absence of density corresponding to the bound LEN. **h**, Comparison of two adjacent CA hexamers within the LEN-bound (blue) and LEN-free (yellow; EMDB: 13423) lattices, showing that LEN binding induces a flattening of the curvature between neighbouring hexamers.

Since the proper cone formation requires precise curvature between neighbouring hexamers, we next quantified the effect of LEN on lattice curvature. To this end, we compared the hexameric CA lattice resolved in our LEN-containing VLPs with published structures of VLPs or virions produced without LEN, as well as to *in vitro*-reconstituted core-like particles (CLPs) ^21,23,34^ (EMDB: 13423, 3465 and 16703), and calculated the interhexamer tilt angles (Fig. 1h, Fig. S4). While the tilt angles between adjacent CA hexamers ranged from 10.7° to 16.9° in previously reported LEN-free structures, the corresponding angle in our LEN-containing VLPs was 6.7°, indicating that LEN binding induces a pronounced flattening of the lattice curvature (Fig. 1h, Fig. S4). To exclude the possibility that the observed flattening was caused by PFO treatment, we also determined the capsid lattice structure of LEN-containing VLPs without PFO perforation and confirmed that a similar flattened lattice conformation was present within intact VLPs (Figs. S5-7, Table S1). Together, these results suggest that LEN binding perturbs the local curvature necessary for the assembly of a functional conical capsid.

To further elucidate how LEN binding alters lattice geometry at the molecular level, we examined the local conformational changes in the protein–protein interfaces that mediate hexamer–hexamer interactions within the lattice (Fig. 2a–c). LEN binding induced structural rearrangements at the CTD–CTD dimer interface, formed by helix 9 ^21–23^ (Fig. 2b), as well as at the trimer interface, formed primarily by helix 10 ^21–23^ (Fig. 2c). Through these changes, LEN disrupts the coordinated lattice geometry required for cone closure, thereby preventing the assembly of a functional conical capsid (Fig. 1). In contrast, the NTD–NTD interface remained unchanged (Fig. 2d), a region that is critical for stabilizing each hexamer through inter-NTD interactions and for coordinating IP6 binding at the central arginine ring. These LEN-induced rearrangements in the interhexamer geometry provide a structural basis for the observed flattening of the lattice curvature in LEN-bound VLPs (Fig. 1h, Fig. S4, Fig. S7). Together, these structural analyses demonstrate that LEN enforces a flattened lattice geometry by inducing specific rearrangements at the CTD–CTD interfaces, thereby reducing the interhexamer curvature and weakening the coordinated architecture necessary to form a functional capsid.

**Fig. 2.**
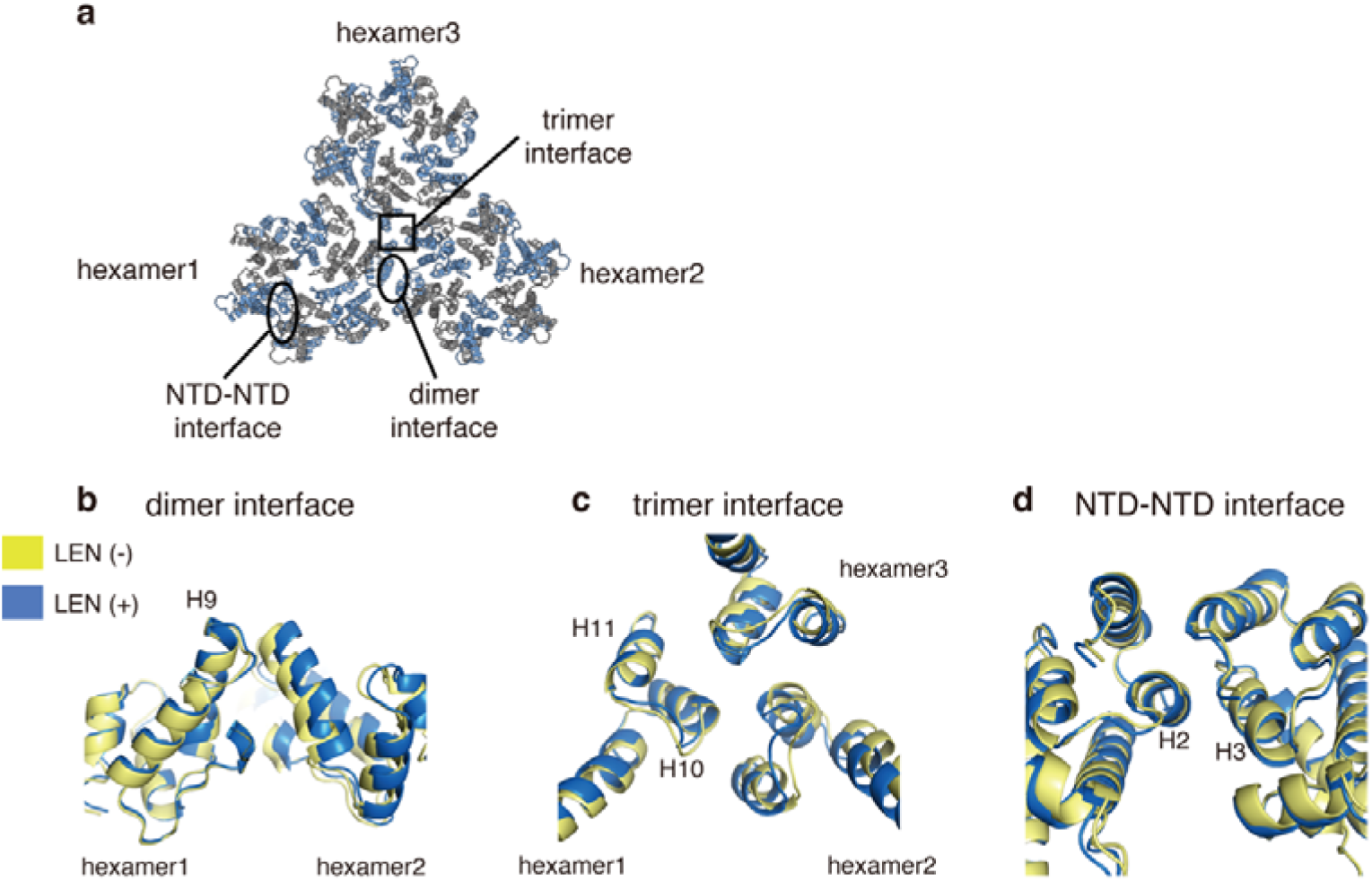
LEN alters the interhexamer interactions in the HIV-1 capsid lattice. **a**, Overview of intra- and interhexamer interfaces within the HIV-1 capsid. **b**, Comparison of the CTD–CTD dimer interface between CA molecules mediated by helix 9, based on the cryo-EM structure of the LEN-bound CA lattice and the cryo-ET structure of a native CA lattice (EMDB: 13423). The LEN-bound CA lattice (blue) was superimposed onto the native CA lattice (yellow) to visualize structural differences at the dimeric interface. Structures were superimposed by aligning one helix 9 of the CA-CTD of hexamer 1. **c**, Comparison of the CTD–CTD trimer interface between CA molecules mediated by primary helix 10, based on the cryo-EM structure of the LEN-bound CA lattice and the cryo-ET structure of a native CA lattice (EMDB: 13423). The LEN-bound CA lattice (blue) was superimposed onto the native CA lattice (yellow) to visualize structural differences at the trimeric interface. Structures were superimposed by aligning one helix 10 of the CA-CTD of hexamer 1. **d**, Comparison of the NTD–NTD interface between CA molecules mediated by helices 2 and 3, based on the cryo-EM structure of the LEN-bound CA lattice and the cryo-ET structure of a native CA lattice (EMDB: 13423). The LEN-bound CA lattice (blue) was superimposed onto the native CA lattice (yellow) to visualize structural differences at the trimeric interface. Structures were superimposed by aligning one helix 3 of the CA-NTD.

### LEN-containing cores exhibit reduced integrity

Given the observed lattice-level perturbations, we next examined whether LEN altered the biochemical properties of the assembled HIV-1 cores. Cores were isolated from HIV-1 virions produced in the presence or absence of LEN using sucrose gradient ultracentrifugation containing IP6. Western blotting for CA demonstrated that cores extracted from the virion produced in the absence of LEN were deposited mainly in fraction 9, whereas those from LEN-containing virions peaked in fraction 10 (Fig. 3a). These results indicate that cores were formed under both conditions; however, the shift to a heavier fraction for the cores produced in the presence of LEN suggests structural alterations, consistent with the cryo-EM images of the VLPs (Fig. 1). In addition, the CA content in the light fractions (1–3), representing either CA dissociated from the lattice during the assay or CA incorporated into virions but not assembled into the lattice, was markedly reduced in the presence of LEN compared with its absence (Fig. 3a). This finding indicates that LEN stabilizes the hexameric lattice of cores, which is consistent with previous reports^2,31^. The peaks of integrase (IN) and viral RNA overlapped with those of the cores under both conditions, indicating that IN and viral RNA were retained within the core in the virion produced in the presence of LEN (Fig. 3b,c). We next assessed the distribution of reverse transcriptase (RT) by measuring RT activity across gradient fractions. Unlike IN and viral RNA, RT was not present in LEN-containing cores, with RT activity in the peak fraction of untreated virions (fraction 9) being approximately 50-fold greater than that in the peak fraction of LEN-treated virions (fraction 10) (Fig. 3g). This marked reduction indicates that RT leaks from LEN-bound capsids. Furthermore, RT leakage from cores was already evident in virions produced in the presence of 2 nM LEN, with similar levels observed at 10 and 50 nM (Fig. S8). To determine whether this LEN-dependent loss of RT is specifically mediated through the FG pocket of CA in the virion, we next tested LEN in HIV-1 bearing the CA M66I mutation, which confers resistance to LEN ^1,3,35^.

**Fig. 3.**
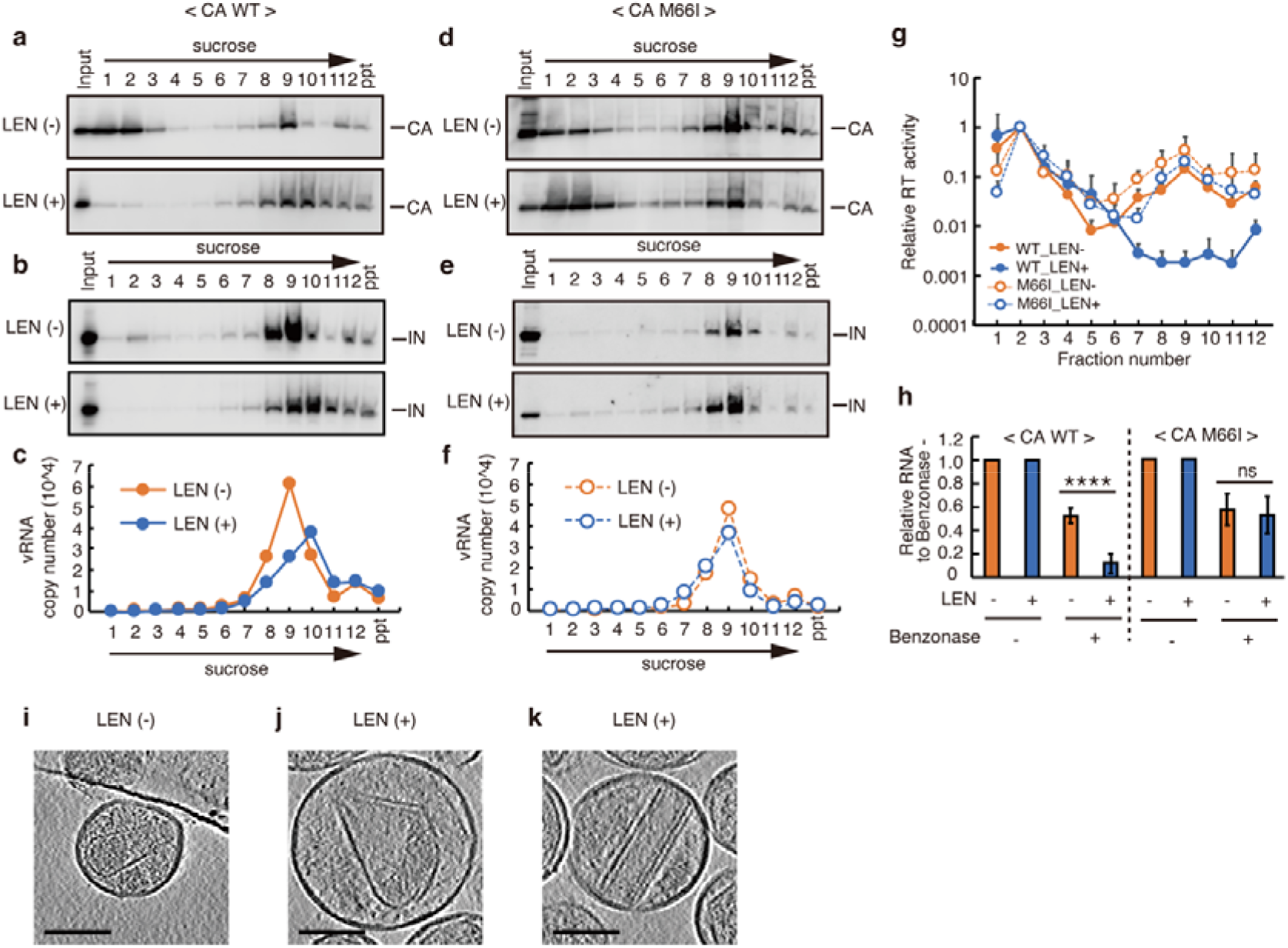
LEN-containing cores in VLPs exhibit reduced integrity and lose reverse transcriptase. **a–c**, Sucrose gradient fractionation of WT HIV-GFP virions produced in the presence or absence of LEN. CA (a) and IN (b) were detected by western blotting. Viral RNA in each fraction was quantified by RT–qPCR (c). Three independent experiments were repeated, and the results were reproduced. **d–f**, Sucrose gradient fractionation of M66I HIV-GFP virions produced in the presence or absence of LEN. CA (d) and IN (e) were detected by western blotting. Viral RNA in each fraction was quantified by RT–qPCR (f). Three independent experiments were repeated, and the results were reproduced. **g**, Reverse transcriptase (RT) activity across sucrose gradient fractions. Red and blue lines indicate WT HIV-GFP produced without or with LEN, respectively; red and blue dashed lines indicate the corresponding M66I mutants produced without or with LEN, respectively. The data are normalized to fraction 2 of each virion sample. The data are presented as the mean ± s.d.; n = 3 independent experiments. **h**, Nuclease digestion assays were performed using cores isolated from the peak sucrose gradient fractions (a–f). Viral RNA levels are shown relative to those in non–benzonase-treated samples. The data are presented as the mean ± s.d.; n = 6 independent experiments. ****P< 0.0001; Student’s t test. **i-k**, Cryo-ET analysis of the Gag VLPs produced in the absence (i) or presence of 50 nM LEN (j,k). Representative tomogram slices of the Gag VLPs produced in the presence of 50 nM LEN (j, k) show partially disrupted cores, including sheet-like and tubular structures with unclosed ends. Scale bars, 100 nm.

Importantly, HIV-1 bearing the CA M66I mutation retained not only IN and viral RNA but also RT within the cores even in the presence of LEN (Fig. 3d–g), demonstrating that RT leakage is specifically due to LEN binding to the FG pocket of CA within the virion. Previous studies have shown that IN binds tightly to viral RNA, serving as a key factor in virion maturation by ensuring proper viral RNA incorporation into the core^36,37^. Furthermore, destabilization of the capsid lattice has been reported to cause loss of RT but not IN^38^. Together, these observations indicate that RT is more susceptible than IN to loss from cores lacking integrity and that the leakage of RT from LEN-containing cores is a clear indicator of compromised capsid integrity.

To further assess the integrity of LEN-containing cores, nuclease digestion assays were performed using cores isolated from the peak sucrose gradient fractions (Fig. 3a–f). These assays revealed that viral RNA was more accessible within LEN-treated cores than in LEN-free cores, whereas M66I cores showed no such change (Fig. 3h). In addition, VLPs produced in the presence of LEN were analysed using cryo–electron tomography (cryo-ET) (Fig. 3i–k; Movies S1–3). Whereas VLPs produced without LEN predominantly contained well-encapsulated cone-shaped capsid (Fig. 3i; Movie S1), those produced in the presence of LEN frequently displayed abnormal structures, including extended sheet-like lattices that failed to curl into closed cones and tubular structures lacking properly sealed ends (Fig. 3j,k; Movies S2,3). These defective structures, in stark contrast to the fully closed conical cores observed without LEN, provide direct structural evidence that LEN impairs the ability of the capsid lattice to assemble into an intact, encapsidated core. Together, these findings demonstrate that LEN binding within the viral core induces a flattened lattice geometry that compromises core integrity, thereby providing a mechanistic basis for its late-phase antiviral activity.

### Structural perturbations impair HIV-1 infectivity

Finally, we sought to determine whether these structural and biochemical perturbations translate to functional defects during HIV-1 replication. First, we investigated the impact of LEN on HIV-1 infectivity using a VSV-G-pseudotyped HIV-GFP virus produced in the presence of LEN. Consistent with previous studies^1,6,31^, the infectivity of HIV-GFP produced in the presence of LEN was reduced to 0.003% in GFP-positive cells, compared with 15% in viruses produced in the absence of LEN, representing ~5,000-fold reduction in infectivity (Fig. 4a). Next, we analysed HIV-1 DNA synthesis, including early reverse transcription (ERT), late reverse transcription (LRT) and integration products, at 9 and 48 hours postinfection (hpi). At both 9 and 48 hpi, the levels of ERT, LRT, and integration products in cells infected with HIV-GFP produced in the presence of LEN were indistinguishable from those in negative control cells infected with virus in the presence of nevirapine, indicating a near complete block of viral DNA synthesis (Fig. 4b–d). In contrast, LEN did not alter the infectivity, HIV-1 DNA synthesis, or integration of HIV-GFP bearing the capsid M66I mutation, which confers resistance to LEN^1,3,35^ (Fig. 4e–h). These results support the idea that the antiviral effect of LEN is attributable to its binding to the FG pocket in the capsid within the virion. Notably, the amount of ERT products was markedly reduced in cells infected with LEN-containing HIV-1. These results support the conclusion that LEN-induced lattice flattening leads to RT loss from the core and thereby prevents efficient initiation of reverse transcription.

**Fig. 4.**
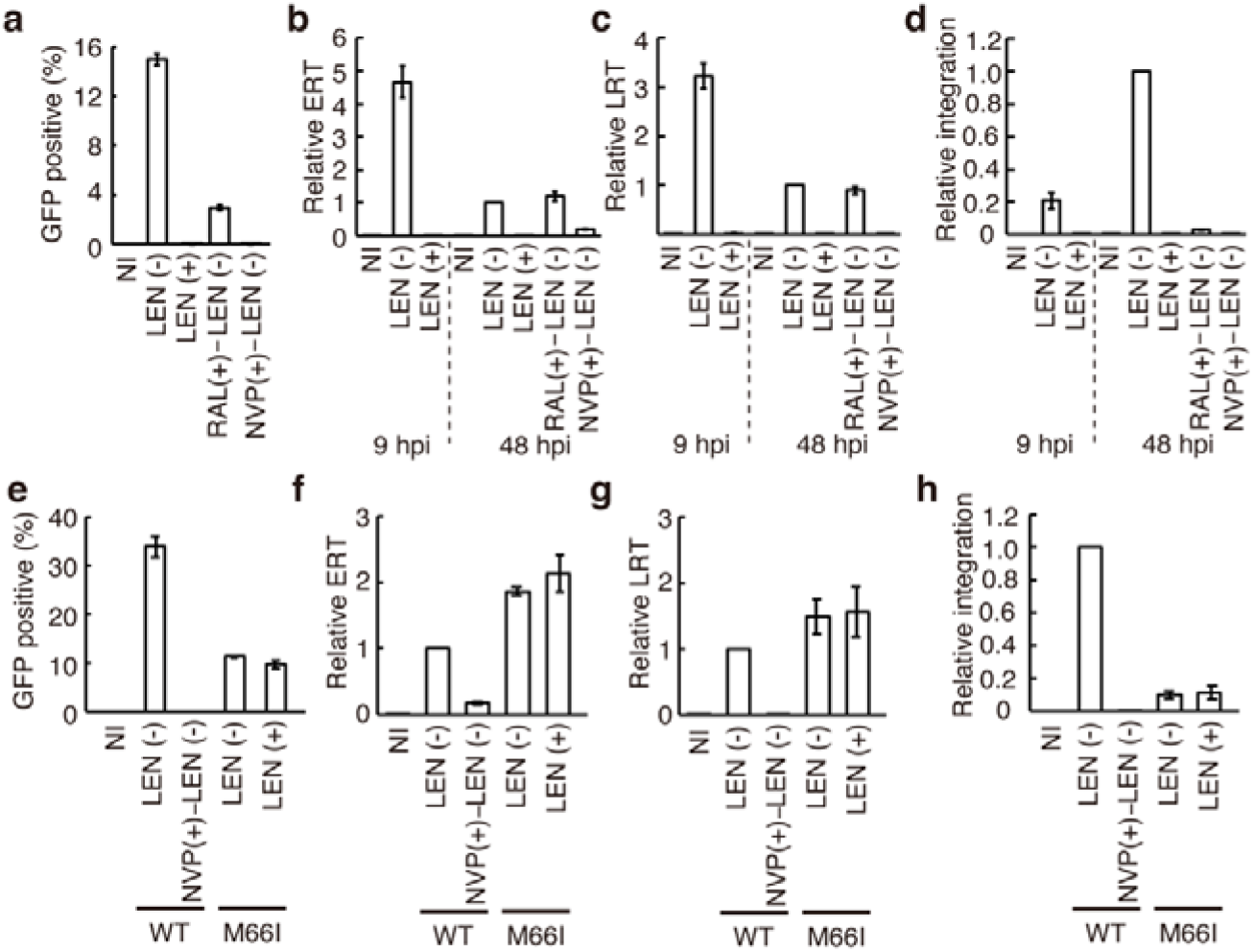
HIV-1 produced in the presence of LEN exhibits reduced infectivity and impaired reverse transcription. **a–d**, Infectivity, reverse transcription (RT) products, and integration in Jurkat cells infected with HIV-GFP virions produced in the presence or absence of 50 nM LEN. Infectivity was quantified by measuring GFP expression at 48 hours postinfection (hpi) (a). Early RT (ERT, b), late RT (LRT, c), and integration (d) products were quantified at 9 or 48 hpi. Raltegravir (RAL, 10 µM) and nevirapine (NVP, 10 µM) were included as controls. **e-h**, Infectivity, RT products, and integration in Jurkat cells infected with HIV-GFP M66I-mutant virions produced in the presence or absence of 50 nM LEN, alongside WT and NVP controls, as shown in the figure. Infectivity was quantified at 48 hpi (e). ERT (f), LRT (g), and integration (h) products were quantified at 48 hpi. The data represent the mean ± s.d. from three independent experiments.

## Discussion

LEN is a first-in-class, long-acting HIV-1 capsid inhibitor and represents one of the most impactful antiviral agents currently in clinical use. Most previous studies have focused on the action of the capsid inhibitor LEN during early postentry steps, where it modulates host factor interactions with the capsid and inhibits reverse transcription, nuclear import, and integration ^1,2,6–8^. In these studies, the potency of LEN has largely been attributed to its ability to disrupt capsid function after infection. However, how LEN perturbs the capsid during the late phase of HIV-1 replication, specifically during virion maturation, has remained unresolved. Here, we addressed this key gap by integrating structural, biochemical, and functional analyses to define how LEN incorporation during virion assembly alters capsid architecture and compromises viral infectivity. Our results reveal that LEN enforces a flattened lattice geometry by inducing rearrangements at the CTD–CTD interfaces, reduces core integrity, and prevents productive reverse transcription and infection. Based on these observations, our data collectively suggest a sequential mechanistic model that links LEN incorporation into assembling virions to subsequent HIV-1 replication failure. Upon incorporation into the maturing virion, LEN binds to the FG pocket at the interface between the CA NTD and CTD, inducing a local rearrangement of the interhexamer geometry and enforcing a flattened lattice conformation. This LEN-induced lattice flattening disrupts the precise spatial organization required for capsid closure, thereby compromising core integrity. As a consequence, cores formed in the presence of LEN prematurely lose RT while still retaining IN and viral RNA and thus fail to support reverse transcription and integration. Therefore, the cascade of structural perturbations triggered by LEN incorporation impairs the HIV-1 replication cycle.

In addition to binding the mature capsid, LEN also interacts with the immature Gag lattice in a distinct manner^1,32^. LEN binding induces conformational changes in the CA region and remodels the architecture of the Gag lattice^32^, which may interfere with the maturation process independently of proteolytic processing of Gag and result in the formation of aberrant capsid structures. Structural analysis of the capsid lattice within VLPs produced in the presence of LEN revealed how LEN disrupts the structure of capsid, demonstrating that LEN binding within the viral core enforces a flattened lattice geometry that compromises capsid integrity. In addition, a previous coarse-grained molecular dynamics study revealed that LEN-treated capsids frequently fail to encapsidate and exhibit impaired maturation^39^. Furthermore, while LEN has been shown to impair the formation of CA pentamers^31,^ which are critical for generating local high curvature and assembling an encapsidated core, our structural analysis indicates that LEN enforces the formation of flattened CA hexamer lattices. Together, these findings suggest that both mechanisms contribute to the production of nonfunctional HIV-1 particles with aberrant capsid structures and thereby inhibit the late steps of HIV-1 replication, providing a mechanistic basis for the profound antiviral efficacy of the clinically approved capsid inhibitor LEN.

Although a pharmacologically relevant concentration of LEN has been reported to be approximately 5 nM^2^, our structural analysis was performed using VLPs produced in the presence of 50 nM LEN (Fig. 1,2) to ensure near-complete lattice occupancy. Notably, the previous study has reported structural abnormalities in HIV-1 virions produced in the presence of 15 nM LEN^1^. Importantly, in our biochemical assays using isolated viral cores, significant RT leakage was observed from cores produced in the presence of 2 nM LEN (Fig. S8). These findings suggest that even partial LEN occupancy, and consequently partial lattice flattening, is sufficient to compromise capsid integrity.

Previous studies using purified mature virion cores showed that LEN disrupts capsid integrity despite stabilizing the capsid lattice^7,8^. These reports further indicate that capsid disruption is non-uniform. Specifically, the narrow ends of viral cores, which contain clustered pentamers and exhibit higher intrinsic curvature, were preferentially destabilized by the treatment of LEN^8^. Although LEN is thought to bind only to CA hexamers and not to pentamers^23,40,41^, rupture predominantly initiated at the narrow ends, suggesting that LEN-induced perturbation of interhexamer geometry preferentially affects high-curvature regions of the lattice. Recent preprint further reported curvature-dependent destabilization of mature HIV-1 capsids by LEN in post-entry cores^42^. Consistent with these observations, our results show that LEN reduces local lattice curvature during virion maturation, perturbing curvature establishment prior to complete cone closure. During viral maturation, these observations support a model in which partial lattice flattening at sub-saturating LEN concentrations is sufficient to impair proper formation of the narrow ends, thereby preventing complete cone closure and productive encapsidation.

Our findings further suggest that capsid curvature is a tunable geometric parameter that can be pharmacologically regulated. Rather than simply hyperstabilizing the capsid lattice, LEN alters interhexamer tilt without disrupting hexamer integrity, thereby biasing assembly toward flattened, non-closable capsid intermediates. This geometric derailment model highlights curvature control as a structural vulnerability of retroviral assembly.

## Methods

### Plasmids

pNL4-3 was obtained from the NIH AIDS Reagent Program. psPAX2 and pMD2. G were gifts from Didier Trono (Addgene plasmid #12260 and #12259). The construct pNL4-3.GFP. R+E-, generated from pNL4-3.Luc.R-E-by replacing the luciferase reporter with GFP and restoring Vpr expression was kindly provided by M. Benkirane.

### Viral production

HIV-1 Env^-^VLPs were produced by transfecting HEK293T Lenti-X cells (Takara/Clontech), confirmed to be mycoplasma-free, with 4 µg of psPAX2 using polyethylenimine (PEI Max) in the presence of 50 nM lenacapavir (LEN; MedChemExpress, HY-111964). VSV-G-pseudotyped HIV-1-GFP was generated by cotransfection of 4 µg of pNL4-3.GFP. R+E- and 1 µg of pMD2-G using PEI Max in the presence of 50 nM LEN. Sixteen hours later, the medium was replaced with DMEM supplemented with 1% FBS supplemented with 50 nM lenacapavir. At 48 hours posttransfection, the medium containing the VLPs was harvested and filtered through a 0.45-µm filter and concentrated by ultracentrifugation through a 20% sucrose cushion. The viral pellet was collected and treated with 100 U/mL DNase (Turbo DNase; Thermo Fisher Scientific) at 37°C for 1 hr. For HIV-1 Env^-^VLPs for cryo-EM analysis, VLPs were further purified by an OptiPrep gradient containing 10, 20, and 30% OptiPrep (Serumwerk Bernburg AG) in STE (10 mM Tris-HCl (pH 7.4), 100 mM NaCl, 1 mM EDTA) from top to bottom. Ultracentrifugation was performed at 45,000 rpm (SW 55Ti rotor) for 2.5 hours at 4°C. The band corresponding to the VLPs was collected and concentrated by ultracentrifugation through an 8% OptiPrep cushion at 42,000 rpm (SW 55Ti rotor) for 1 hour at 4°C.

### Quantification of infectivity and viral DNA

Jurkat cells, confirmed to be mycoplasma-free, were infected with VSV-G-pseudotyped HIV-1-GFP produced in the presence or absence of LEN. The multiplicity of infection (MOI) was set to 0.3, as determined by the percentage of GFP-positive Jurkat cells infected with VSV-G-pseudotyped HIV-1-GFP produced in the absence of LEN. For infections with viruses produced in the presence of LEN, equivalent amounts of viral RNA to that of LEN-free virus were used. After 3 hours, the cells were washed twice with PBS, and the culture medium was replaced. At 48 hours postinfection (hpi), the cells were collected and fixed with 4% paraformaldehyde and subjected to flow cytometry analysis using a BD FACSVerse (BD Biosciences). For viral DNA quantification, at 9 or 48 hpi, DNA was extracted from infected cells using a NucleoSpin Tissue Kit (Takara/Clontech) and subjected to qPCR. For the quantification of ERT and LRT products, 12.5 ng of DNA (quantified using a Qubit) was used in each reaction^43–45^. Integrated HIV DNA was quantified by the Alu–PCR method as described previously ^44,46,47^, using 12.5 ng of DNA (quantified using a Qubit). ERT, LRT and integrated HIV DNA concentrations were normalized to b-globin concentrations. The primers used for viral DNA quantification are listed in Table S2.

### Sucrose gradient fractionation of HIV-1 cores

VSV-G pseudotyped HIV-1-GFP particles produced in the presence or absence of LEN were subjected to sucrose gradient fractionation as previously described, with slight modifications^8^. Viral input was normalized across samples on the basis of viral RNA levels and quantified by qPCR using primers specific for the LRT product prior to loading. Virions were subjected to ultracentrifugation through a 1% Triton X-100 layer into a linear 30–70% sucrose density gradient containing 10 ml of 10 mM HEPES (pH 7.5), 150 mM NaCl, and 200 μM IP6 (Sigma). The samples were applied to the top of the gradient and centrifuged at 27,000 rpm for 16 hours at 4 °C using a Beckman SW40Ti rotor. Following centrifugation, 1 ml fractions were collected sequentially from the top of the gradient. CA and IN were detected in each fraction by western blotting using serum or antibodies against CA (NIH AIDS Reagent Program, Cat# 4250) and IN (Santa Cruz, sc-69721). For quantification of viral RNA in each fraction, RNA was extracted using the NucleoSpin RNA Virus Kit (Takara/Clontech). Reverse transcription was then performed, followed by qPCR using primers specific for LRT. RT activity in each fraction was quantified using a qPCR-based product-enhanced reverse transcriptase (PERT) assay, as previously described^48,49^.

### Nuclease sensitivity assay

Five microlitres of the peak fraction from the sucrose gradient was diluted fivefold with 20 µl of sucrose gradient buffer lacking sucrose and treated with 50 U/ml benzonase (Merck Millipore) in the presence of 5 mM MgCl□. After incubation at 37 °C for 30 min, the reaction was stopped by the addition of 10 mM EDTA. Viral RNA was then purified using the NucleoSpin RNA Virus Kit (Takara/Clontech). Reverse transcription was performed, followed by qPCR using primers specific for LRT.

### Cryo-EM structure determination

#### Cryo-EM sample preparation

To prepare the VLPs treated with PFO, the VLPs were incubated with 0.1 mg/ml PFO (CUSABIO TECHNOLOGY LLC) and 200 μM IP6 (Sigma) at room temperature for 2 hours. For the preparation of the cryo-EM specimen, Quantifoil grids (R2/2, 200-mesh, copper; Quantifoil Micro Tools GmbH) were pretreated with acetone and glow-discharged with a PIB-10 system (Vacuum Device). Two-microlitre samples were applied to the grids, and the grids were then blotted at 100% humidity and 18 °C in a Vitrobot Mark IV System (Thermo Fisher Scientific) or EM GP2 (Leica Microsystems). The blotted grids were immediately plunged into liquid ethane.

#### CryoEM SPA data collection

Cryo-EM images for SPA were acquired on a Krios G4 (Thermo Fisher Scientific) transmission electron microscope operated at 300 kV with a K3 direct electron detector (Gatan) with a GIF-Biocontinuum energy filter (Gatan) with a 20-eV slit width. Automated data collection was performed with EPU software (Thermo Fisher Scientific) with a nominal magnification of 81,000, defocus values between −0.8 and −2.4 µm, and an electron dose of 1.01–1.03 e-/Å2 per frame over 50 frames.

#### Cryo-EM SPA image processing

Image processing was performed with cryoSPARC^50^. Micrograph movie frames were aligned, dose weighted, and CTF estimated using patch motion correction and patch CTF in cryoSPARC. The particle images were automatically picked with a Blob picker. Junk particles were then removed by 2D classification. For the first round of 3D heterogeneous refinement, the structure of the intact mature CA hexamer (EMDB: 3465) and decoy maps derived from a 100 Å lowpass-filtered CA hexamer map (EMDB: 3465) were used as the initial 3D model. Iterative rounds of heterogeneous refinement and 2D classification were performed to remove junk particles. Then, particles exposing the capsid from ruptured virions were manually excluded. The selected particles were subjected to reference-based motion correction, global CTF refinement, local CTF refinement, and non-uniform refinement. The global resolutions were based on the gold-standard Fourier shell correlation curve (FSC=0.143) criteria. The workflows for data processing are shown in Fig. S3 (the LEN-bound HIV-1 capsid lattice within VLPs treated with PFO) and Fig. S6 (the LEN-bound HIV-1 capsid lattice within intact VLPs).

#### Model building and refinement

The initial atomic models of HIV-1 CA (PDB: 4XFX) were rigid bodies fitted to the cryo-EM map via UCSF Chimera and ChimeraX ^51,52^. The models were subjected to automatic refinement with phenix.real_space_refine^53^ and manually edited with Coot ^54^. The models were validated with the MolProbity tool^55^.

#### CryoET data collection

Cryo-EM images for tomography were acquired on a Krios G4 (Thermo Fisher Scientific) transmission electron microscope operated at 300 kV with a K3 direct electron detector (Gatan) and a GIF-Biocontinuum energy filter (Gatan) with a 20-eV slit width. Each micrograph was collected at a nominal magnification of 26,000 (pixel size of 3.37 Å), with defocus values between −6 and −8 µm and an electron dose of 2.53 e-/Å2. Tilt-series data collection was performed with a dose-symmetric scheme starting from 0° with a 3° tilt increment and an angular range of ±60° using Tomography 5 software (Thermo Fisher Scientific).

#### CryoET data processing

The micrographs were processed using the AreTomo2 software^56^. The tomograms were binned 4x and reconstructed with weighted back projection. Reconstructed tomograms were also imported and improved using IsoNet software^57^. Reconstructed volumes were visualized with IMOD package^51,52,58^.

#### Use of large language models

GPT-4o and ChatGPT5 were used for grammatical correction of the text. No original sentence was produced by AI.

## Data availability

Cryo-EM maps have been deposited in the Electron Microscopy Data Bank (EMDB) under accession numbers EMD-67115 (LEN-bound HIV-1 capsid lattice within VLPs treated with PFO) and EMD-67116 (LEN-bound HIV-1 capsid lattice within intact VLPs). The coordinates have been deposited in the Protein Data Bank (PDB) under accession numbers 9XQG (LEN-bound HIV-1 capsid lattice within VLPs treated with PFO) and 9XQH (LEN-bound HIV-1 capsid lattice within intact VLPs).

## Acknowledgements

We wish to thank S. P. Goff and M. Benkirane for the pNL4-3.Luc.R-E- and pNL4-3.GFP. R+E-molecular clones. This work was supported by the Japan Agency for Medical Research and Development (AMED) (JP25fk0410078, JP25fk0310547, JP25fk0310530, JP25fk0310527), the JSPS KAKENHI (JP25K02504), the National Institute of Global Health and Medicine (22T003 and 23A1017), the Kato Memorial Bioscience Foundation, the Mitsubishi Foundation, and the Takeda Science Foundation (all to SM). HT was supported by the National Institute of Global Health and Medicine (25A1014). KM was supported by the Japan Agency for Medical Research and Development (AMED) (JP22ama121037 and JP243fa627005), the JSPS KAKENHI (JP20H05873), the National Institute of Global Health and Medicine (23A1017), and the Takeda Science Foundation.

## Author contributions

Conceptualization: H.T. and S.M.; Methodology: H.T., T.O., S.K., K.M., and S.M.; Investigation: H.T., R.M., T.O., N.S., and S.M.; Writing – Original Draft: H.T. and S.M.; Writing – Review & Editing: H.T., R.M., T.O., S.K., K.M., and S.M.; Funding Acquisition: H. T, K.M., and S.M.; Resources: S.M.; Supervision: S.M.

## Competing interests

The authors declare that they have no competing interests.

**Correspondence and requests for materials** should be addressed to Shinichi Machida.

## Notes

### Competing Interest Statement

The authors have declared no competing interest.

