## Supplementary Information for "Structural basis of lenacapavir-induced HIV-1 capsid disruption during virion maturation"

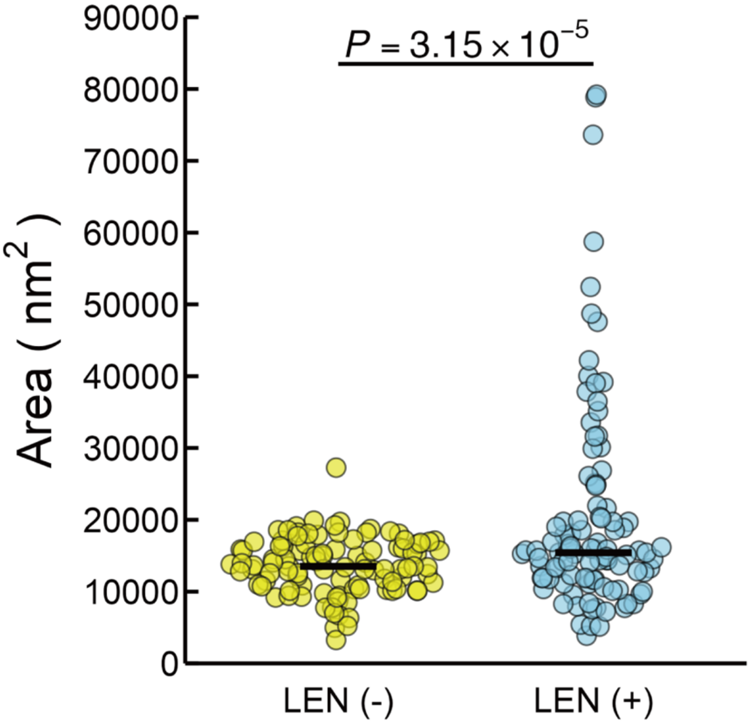

**Fig. S1. Size distribution of HIV-1 VLPs produced in the presence or absence of LEN.**

Scatter plot showing the area of the virus-like particles (VLPs) produced from 293T cells in the absence (yellow) or presence (blue) of 50 nM LEN, as measured from cryo-EM micrographs. Each dot represents an individual VLP (*n* = 101 per group), and the horizontal bars indicate the median. Compared with the untreated controls, the VLPs produced in the presence of LEN had a significantly larger particle area ($P = 3.15 \times 10^{-5}$, unpaired two-tailed Welch's t test).

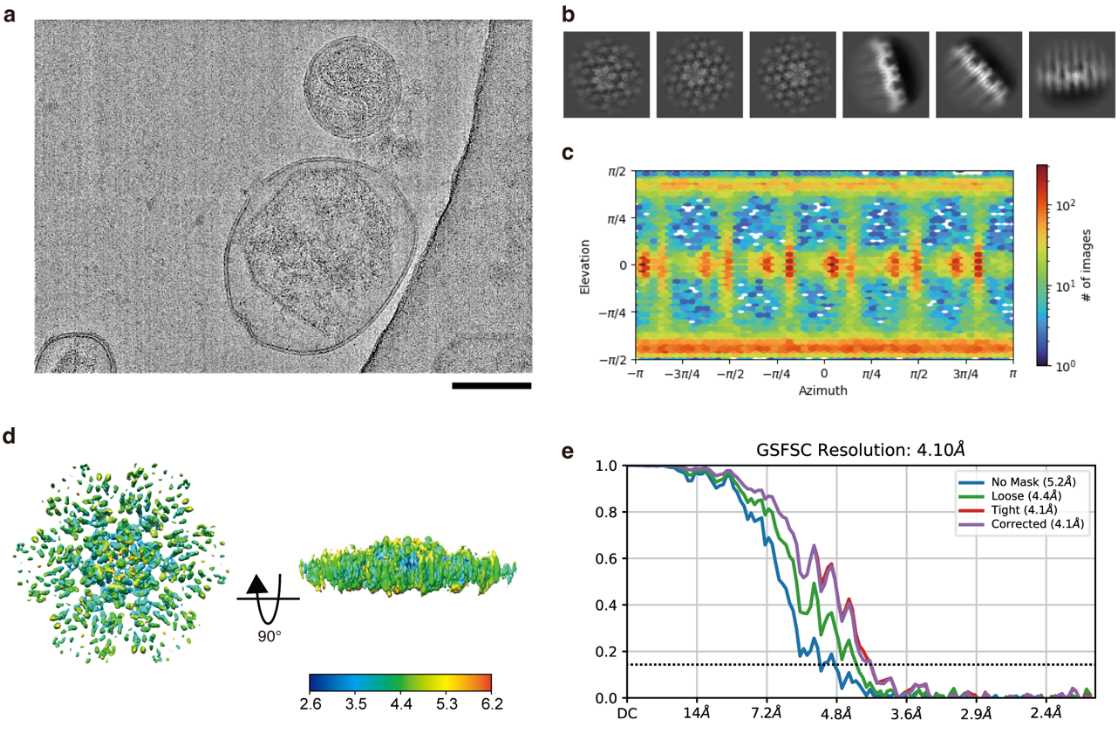

**Fig. S2. Cryo-EM analysis of the LEN-bound HIV-1 capsid lattice within VLPs treated with PFO.**

**a**, Representative micrograph from the cryo-EM imaging dataset. Scale bar, 100 nm. **b**, Selected 2D class averages. **c**, Euler angle distribution plots of the particles of the LEN-bound CA lattice contributing to the final reconstruction. **d**, Local resolution map of the LEN-bound CA lattice. **e**, Fourier shell correlation (FSC) curve for the LEN-bound CA lattice. The dashed line indicates the FSC threshold of 0.143.

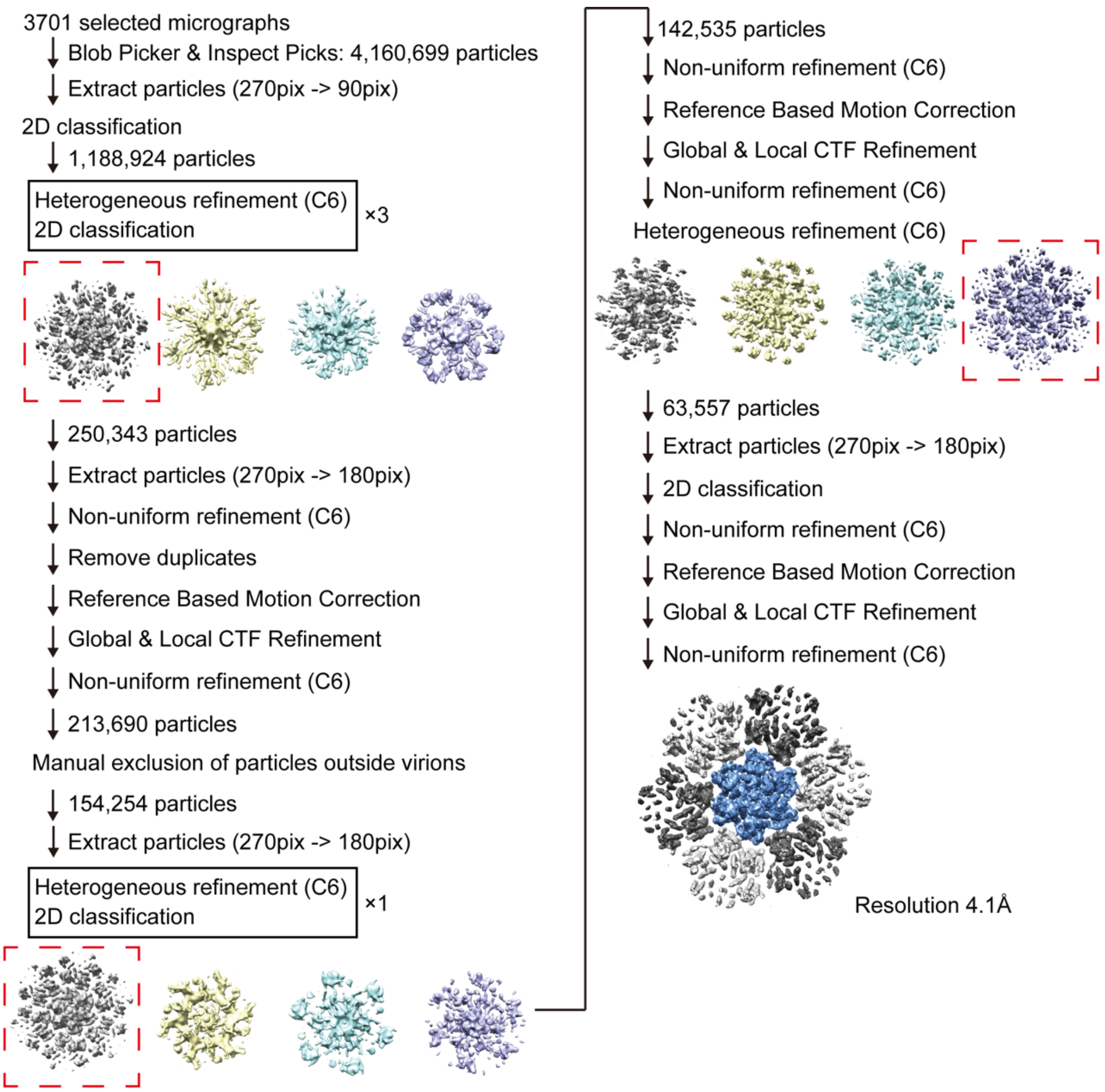

**Fig. S3. Data processing workflow for the LEN-bound HIV-1 capsid lattice within VLPs treated with PFO.**

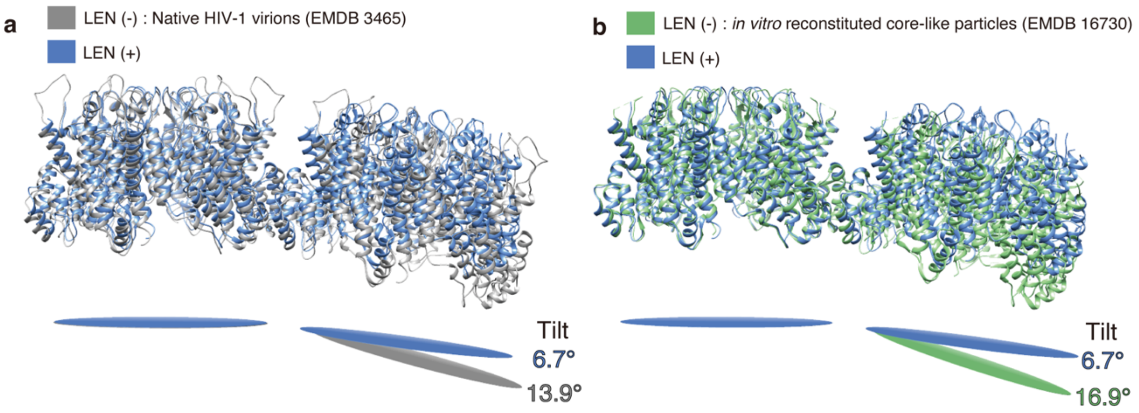

**Fig. S4. Comparison of interhexamer angles between LEN-bound CA lattices and previously reported LEN-free structures.**

**a.** Comparison of two adjacent CA hexamers within the LEN-bound lattice (blue) and the native LEN-free lattice of HIV-1 virions produced without LEN (grey; EMDB: 3465).

**b.** Comparison of two adjacent CA hexamers within the LEN-bound lattice (blue) and the LEN-free lattice of *in vitro* reconstituted core-like particle (green; EMDB: 16703) lattices.

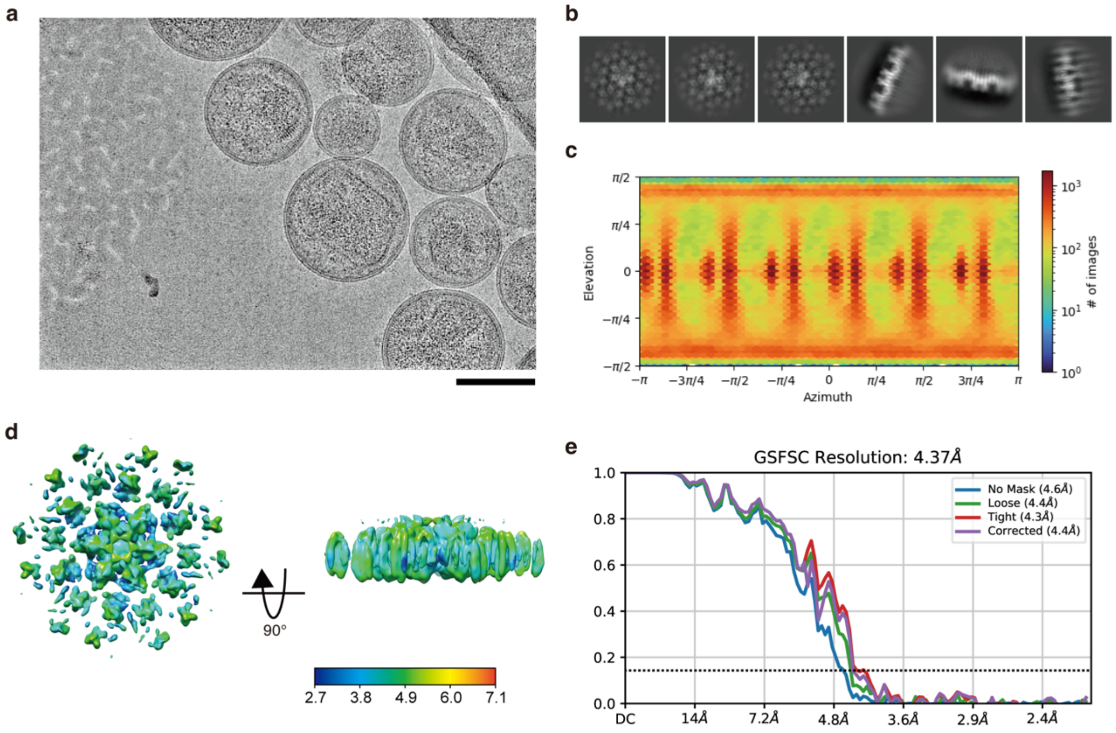

**Fig. S5. Cryo-EM analysis of the LEN-bound HIV-1 capsid lattice within intact VLPs.**

**a**, Representative micrograph from the cryo-EM imaging dataset. Scale bar, 100 nm. **b**, Selected 2D class averages. **c**, Euler angle distribution plots of the particles of the LEN-bound CA lattice contributing to the final reconstruction. **d**, Local resolution map of the LEN-bound CA lattice. **e**, Fourier shell correlation (FSC) curve for the LEN-bound CA lattice. The dashed line indicates the FSC threshold of 0.143.

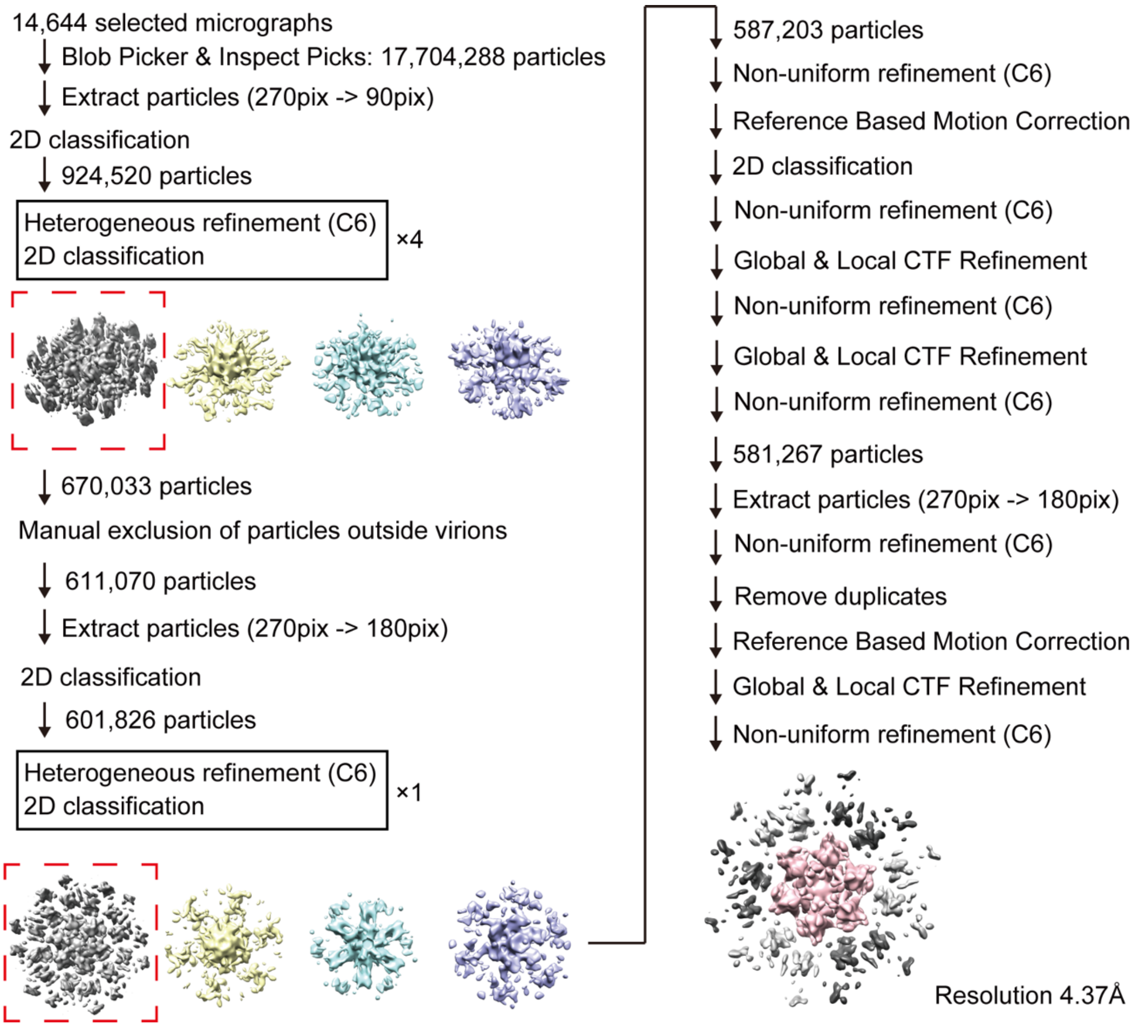

**Fig. S6. Data processing workflow for the LEN-bound HIV-1 capsid lattice within intact VLPs.**

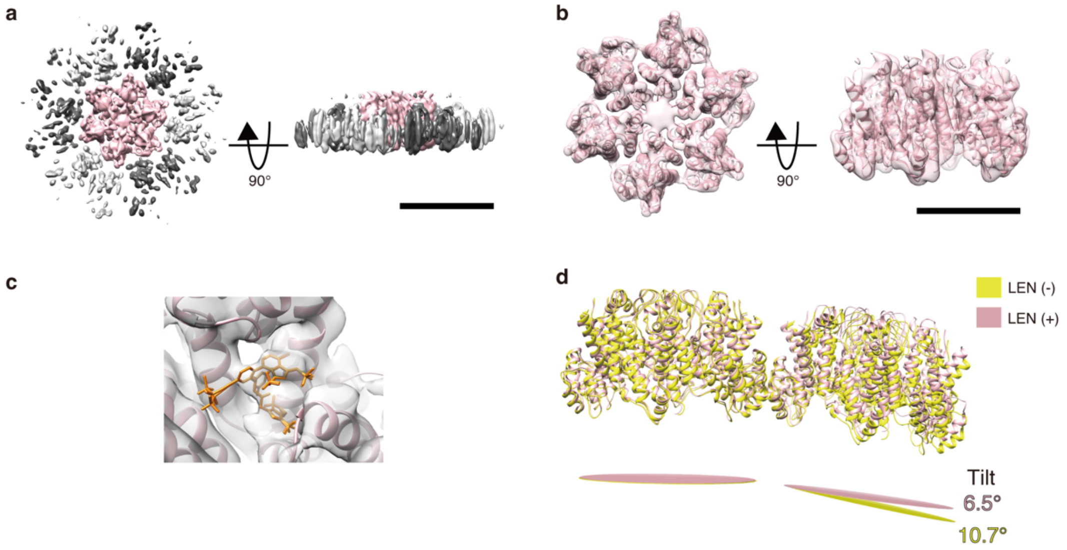

**Fig. S7. LEN alters the interhexamer interactions in LEN-containing VLPs within intact VLPs.**

**a**, Cryo-EM density map of the CA lattice within LEN-containing VLPs, shown from the top (left) and side (right) views. Scale bar, 100 Å.

**b**, The atomic model of a CA hexamer in the LEN-bound lattice fitted into the cryo-EM density map shown in (a). Scale bar, 50 Å.

**c**, Close-up view of the cryo-EM map of a LEN-bound CA lattice and the atomic model showing the density corresponding to a bound LEN.

**d**, Comparison of two adjacent CA hexamers within the LEN-bound (grey) and LEN-free (yellow; EMDB: 13423) lattices, showing that LEN binding induces a flattening of the curvature between neighbouring hexamers.

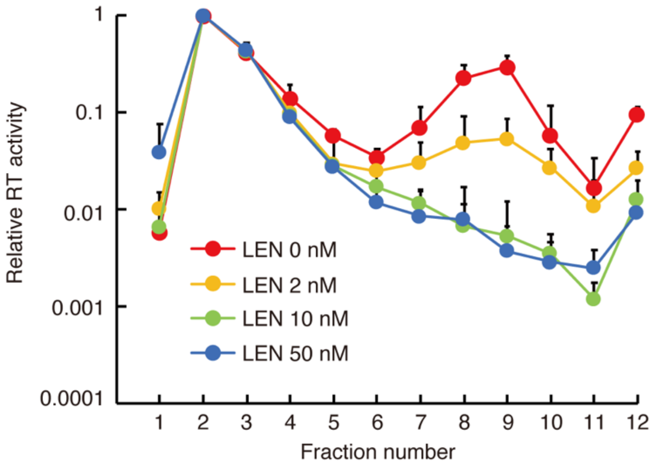

**Fig. S8. Effect of LEN concentration on the leakage of reverse transcriptase from HIV-1 cores.**

Reverse transcriptase (RT) activity across sucrose gradient fractions. VSV-G-pseudotyped HIV-1-GFP was produced by cotransfection of pNL4-3.GFP. R+E- and pMD2-G using PEI Max in the absence or presence of LEN (2, 10, or 50 nM). At 48 hours post-transfection, the medium containing the VLPs was harvested and filtered through a 0.45-µm filter, treated with 100 U/mL DNase, and subjected to sucrose gradient fractionation. Viral input was normalized across samples on the basis of viral RNA levels. Red, yellow, green, and blue lines represent WT HIV-GFP produced without LEN or with LEN (2, 10, or 50 nM), respectively; the data are normalized to fraction 2 of each virion sample and are shown as the mean ± s.d. (n = 3 independent experiments).

### Table S1. Cryo-EM data collection, refinement and validation statistics

| | #1 LEN-bound CA lattice within VLPs treated with PFO (EMD-67115) (PDB ID: 9XQG) | #2 LEN-bound CA lattice within VLPs (EMD-67116) (PDB ID: 9XQH) |
|---|---|---|
| **Data collection and processing** | | |
| Magnification | 81,000 | 81,000 |
| Voltage (kV) | 300 | 300 |
| Electron exposure (e–/Å$^2$) | 51.7 | 50.8 |
| Defocus range (μm) | -0.8 to -2.4 | -0.8 to -2.4 |
| Pixel size (Å) | 1.06 | 1.06 |
| Initial particle images (no.) | 4,160,699 | 17,704,288 |
| Final particle images (no.) | 63,097 | 565,760 |
| Map resolution (Å) | 4.1 | 4.37 |
| FSC threshold | (FSC=0.143) | (FSC=0.143) |
| **Refinement** | | |
| Initial model used (PDB code) | PDB: 4XFX | PDB: 4XFX |
| Model composition | | |
| Non-hydrogen atoms | 4992 | 4992 |
| Protein residues | 1248 | 1248 |
| Ligands | 0 | 0 |
| R.m.s. deviations | | |
| Bond lengths (Å) | 0.002 | 0.005 |
| Bond angles (°) | 1.125 | 1.143 |
| Validation | | |
| MolProbity score | 0.63 | 0.89 |
| Clash score | 0 | 0.95 |
| Poor rotamers (%) | 0 | 0 |
| Ramachandran plot | | |
| Favored (%) | 97.30 | 97.47 |
| Allowed (%) | 2.70 | 2.53 |
| Disallowed (%) | 0 | 0 |

**Table S2. Primers used for quantification of HIV DNA**

| Name | Sequence (5'-3') | Target | Reference |
|---|---|---|---|
| ERT_F | GGCTAACTAGGGAACCCACTG C | Early RT product | |
| ERT_R | CAACAGACGGGCACACACTAC T | Early RT product | |
| LRT_F | TGTGTGCCCGTCTGTTGTGT | Late RT product | ([45]) |
| LRT_R | CTTCAGCAAGCCGAGTCCTG | Late RT product | ([45]) |
| L-M667 | ATGCCACGTAAGCGAAACTCT GGCTAACTAGGGAACCCACTG | Integrated HIV (Alu-PCR (1st round PCR)) | ([44,46,47]) |
| Alu 1 | TCCCAGCTACTGGGGAGGCT GAGG | Integrated HIV (Alu-PCR (1st round PCR)) | ([44,46,47]) |
| Alu 2 | GCCTCCCAAAGTGCTGGGATT ACAG | Integrated HIV (Alu-PCR (1st round PCR)) | ([44,46,47]) |
| Lambda T | ATGCCACGTAAGCGAAACT | Integrated HIV (Alu-PCR (2nd round PCR)) | ([44,46,47]) |
| AA55M | GCTAGAGATTTTCCACACTGA CTAA | Integrated HIV (Alu-PCR (2nd round PCR)) | ([44,46,47]) |
| β-globin-F | CCCTTGGACCCAGAGGTTCT | β-globin | ([44,46,47]) |
| β-globin-R | CGAGCACTTTCTTGCCATGA | β-globin | ([44,46,47]) |

**Movie S1-S3. Cryo-electron tomography of HIV-1 VLPs produced in the presence or absence of LEN.**

Movie S1. Cryo-ET of VLPs produced in the absence of LEN, corresponding to Fig. 3i.

Movie S2. Cryo-ET of VLPs produced in the presence of LEN, showing aberrant sheet-like lattices, corresponding to Fig. 3j.

Movie S3. Cryo-ET of VLPs produced in the presence of LEN, showing tubular or incompletely closed capsid structures, corresponding to Fig. 3k.
